# An empirical, 21st century evaluation of phrenology

**DOI:** 10.1101/243089

**Authors:** O. Parker Jones, F. Alfaro-Almagro, S. Jbabdi

## Abstract

Phrenology was a nineteenth century endeavour to link personality traits with scalp morphology, which has been both influential and fiercely criticised, not least because of the assumption that scalp morphology can be informative of underlying brain function. Here we test the idea empirically rather than dismissing it out of hand. Whereas nineteenth century phrenologists had access to coarse measurement tools (digital technology referring then to fingers), we were able to re-examine phrenology using 21^st^ century methods and thousands of subjects drawn from the largest neuroimaging study to date. High-quality structural MRI was used to quantify local scalp curvature. The resulting curvature statistics were compared against lifestyle measures acquired from the same cohort of subjects, being careful to match a subset of lifestyle measures to phrenological ideas of brain organisation, in an effort to evoke the character of Victorian times. The results represent the most rigorous evaluation of phrenological claims to date.

## Introduction

According to Franz Joseph Gall, the founder of phrenology, those of a *mirthful* disposition (i.e. those who like to laugh) should expect to find two prominent bumps on the forehead when compared to their more dour contemporaries^1^. For nearly two centuries now, the academic community has openly mocked phrenology, yet the approach has seen moments of near redemption. In 1998 for example, electrical stimulation of the pre-SMA, a brain area near the “mirth” bump described by Gall, reportedly caused a patient to laugh^2^. More likely than not, Gall's association of this area with an “Organ of Mirthfullness” was accidental. It nonetheless frames the question empirically: does the local shape of the head reflect aspects of individual psychology?

A good reason to be sceptical about this is that the methodology behind phrenology was dubious even by the standards of the early 19^th^ century. Phrenologists asserted the location of an “Organ of Amativeness” (describing “the faculty that gives rise to sexual feeling”), for example, by probing the heads of “emotional” young women as well as the recently widowed; they further hypothesised the location for an “Organ of Combativeness” by, inversely, searching for flat regions on the scalps of peaceable “Hindoos and Ceylonese”^3^ (p. 46). The phrenological approach therefore relied on tenuous and perhaps even offensive stereotypes about different social groups. Gall's science of “bump reading” would ultimately be abandoned as much for its fixation on social categories as for an inability within the scientific community to replicate its findings. It was these scientific failings that would be exposed by anatomists like Paul Broca and Carl Wernicke, who pioneered the alternative neuroscientific method of lesion-symptom mapping^4,5^. Whereas lesion-symptom mapping described the brain directly, phrenology had to assume that scalp morphology correlated with local brain function indirectly. Even more damning: the results of lesion-symptom mapping contradicted those of phrenology. For instance, Broca and Wernicke identified lateral language areas in cortex roughly around the ear, where later phrenologists had asserted that the “Organ of Language” could be found below the eye^1^. In retrospect, the phrenological proposition that the brain is organised around functionally discrete modules was prescient. However, the idea that the brain's soft tissue might exert a significant effect on skull shape was, and is, nonsense. *Or is it?*

In this study, we sought to test the 19^th^ century claims of phrenology by using 21^st^ century scientific methods. We asked whether local changes in scalp morphology, measured reliably in almost six thousand subjects, do or do not correlate with the “faculties” that Gall described. For historical completeness, we also asked a second question: does local scalp morphology reflect the brain's underlying morphology? We asked this question because phrenologists believed that inspecting the outer surface of the head provided an indirect measure of brain shape based on the assumption that the softness of the skull during development should allow it to yield under the pressure of locally expanding cortical structures^6-8^. For data, we turned to the world's largest brain-imaging study, currently acquiring MRI and other data for 100,000 subjects^9,10^. We used all of the data from the first public release (5,724 subjects). The original scans were separated into parts representing the brain and parts representing the outer surface of the head; and we focused on the outer surface of the head. By applying modern methods from neuroimaging—such as registration and normalisation, random field theory and mass univariate analysis—to the study of the cranium, we were able to search for statistical relationships between local head shape and lifestyle measures, which we took to reflect the “faculties” (or in modern terms “functions”) associated with phrenology. Although we did not expect to find any significant effects between lifestyle measures and head shape, we do believe it is important for scientists to test ideas, even unfashionable or offensive ones, and not be content dismissing them out of hand. This study therefore represents the most rigorous evaluation of phrenological claims ever attempted and aims to offer either vindication or the strongest objection yet against phrenology.

## Methods

### Data

We used anatomical brain-imaging data sampled from the UK Biobank Imaging study (http://imaging.ukbiobank.ac.uk). These data are representative of the largest neuroimaging study to date, aiming to acquire MRI and personal measures (including questionnaires and cognitive tests) for 100,000 subjects^9,10^. We used all of the available data from the first public release of 5,724 subjects (2,693 male, aged 45 to 78 years; mean=62 years, standard deviation=7 years; see Supplementary Figure 1).

**Figure 1:**
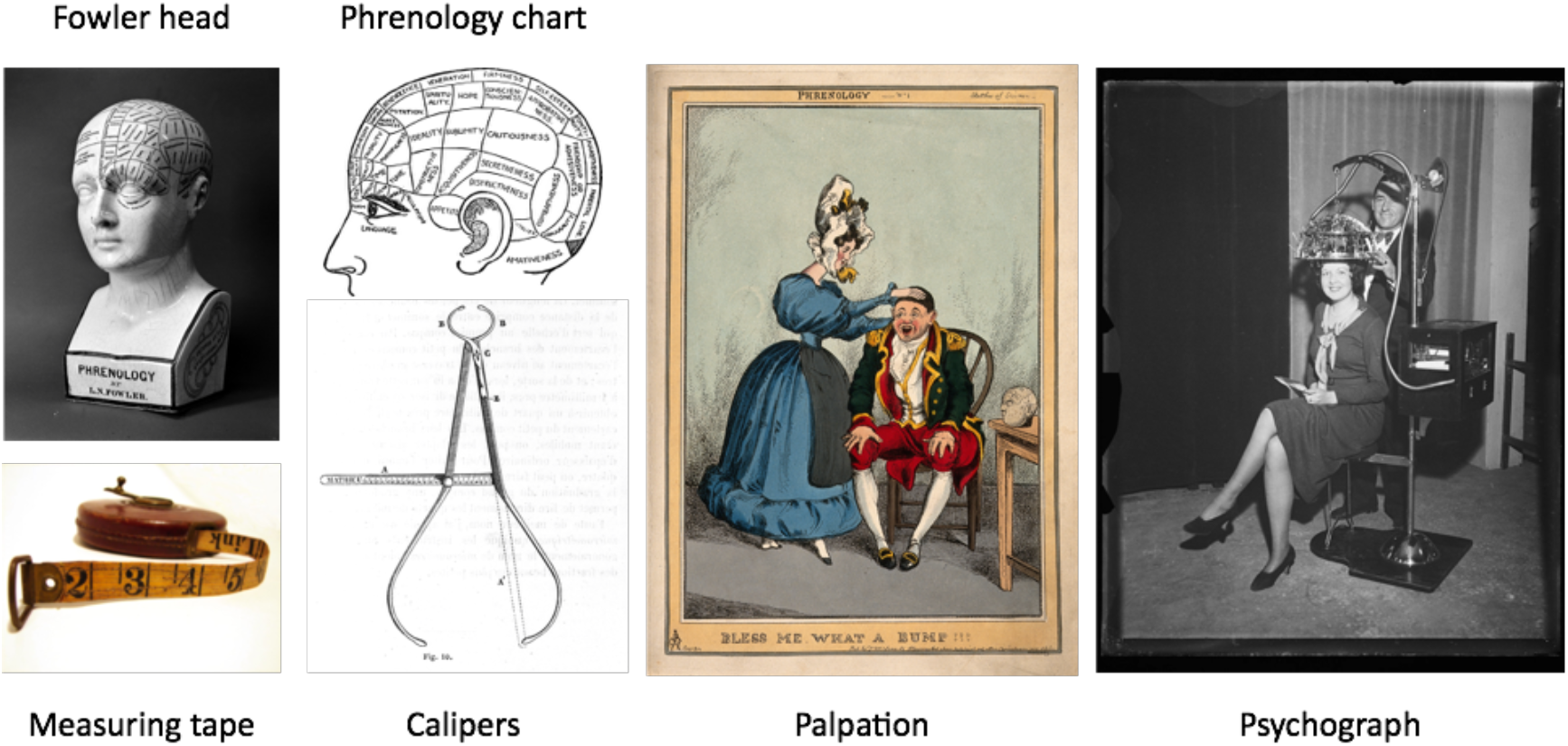
Traditional tools of phrenology: Fowler head^11^, Phrenology chart^12^, measuring tape^13^,^14^, palpation^15^, psychograph^16^. calipers:

### Pre-processing

Each subject's T1-weighted structural scan was processed using the FSL Scalp Extraction Tool (SET)^17,18^. SET is used to produce an estimate of both inner and outer surfaces of the head (Figure 2). Neuroimaging studies typically retain the extracted brain. We discarded the brain to focus instead on the scalp surface.

**Figure 2:**
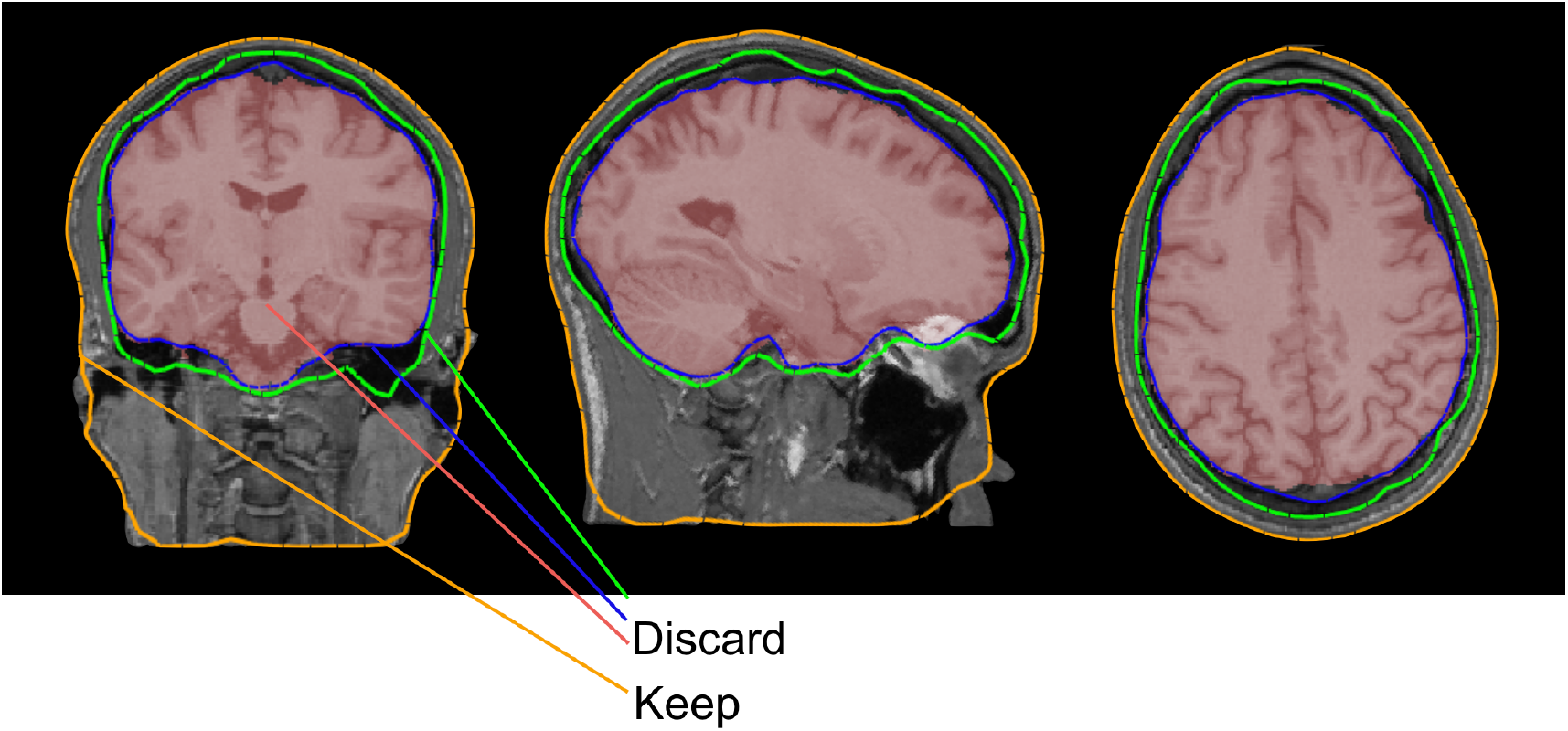
Schematic of FSL’s scalp extraction tool, which identifies various tissue boundaries (red=brain; blue=pial boundary; green=inner skull surface; orange=outer scalp surface). Phrenology is focused on the scalp (outer surface of the head).

T1-weighted images were linearly aligned to a standard brain template (MNI152) using FLIRT^19,20^, and the same transformations were applied to the vertex coordinates of the scalp surfaces of each subject. All scalp surfaces for all subjects were therefore aligned with one-to-one correspondence between the vertices, making it possible to compare scalps between subjects within a common space. In addition, we applied a hand-drawn mask to exclude surface vertices below the nose, as these exhibited high levels of between-subject variation and were typically excluded by phrenologists (e.g. note the grey regions in Figure 5).

Although some phrenologists took global measures of the head using a measuring tape or calipers (Figure 1), this practice was not unique. What was unique to phrenology was its emphasis on local head curvature, or “bumps”, making it our focus here. We calculated the mean (signed) curvature at each vertex of each individual surface projection^21^. This gave us 40,962 vertex measures per subject (see Figure 4, panel A) which we then compared against a set of lifestyle measures drawn from the same subjects.

### Lifestyle measures and phrenological faculties

Philosophically, phrenology was organised around the metaphor of the brain as a collection of physical “organs” with identifiable functions, such as “language” or “love”, or an “impulse to propagation”. In phrenology, these functions are referred to as “faculties”. Although these faculties diverge from the familiar functions mapped by neuroimaging in the 20^th^ and 21^st^ centuries, in this regard the approaches do not differ in kind^22^.

In addition to MRI, the UK Biobank Imaging study includes data from numerous questionnaires and cognitive tests, which we refer to collectively as “lifestyle measures”. Subject responses to these lifestyle measures could be binary (“Do you live with your parents?”) or integer-valued (“How many sexual partners have you had?”). Some integer-valued responses required closed-set answers (for example, “How often do you eat beef?” given a range of options from 0-4, where 0 means “never” and 4 means “I eat beef daily”). We used the lifestyle measures as proxies for 23 common phrenological “faculties”^1^.

Gall originally proposed 27 faculties^23,24^. From these, we selected a subset of 23 faculties for which we found compelling lifestyle measures in the UK Biobank. To illustrate, we associated the faculty of combativeness (argumentativeness) with lawyers, and we associated cunning with scientists. By connecting the faculty of “cunning” to our own profession, we are following a phrenological tradition which is evident for example in Fowler and Fowler's^25^ choice to cite Gall's skull as an example of “Causality” (also referred to as “metaphysical perspicuity”—intended, it seems, to be a good thing).

We give the full list of Faculties and associated lifestyle measures in Table 1, noting that: letter fluency (Faculty XIV) is the number of words starting with the letter “s” that the subject could produce in one minute and concept interpolation (Faculty XX) is a fluid-intelligence test which records one's capacity to solve problems that require logic and reasoning independent of acquired knowledge (where each subject had 2 minutes to complete as many questions as possible from the test). Four of Gall's Faculties were excluded from our study because we could not find appealing proxies in the set of lifestyle measures. The excluded Faculties were: XIII (recollection for persons), XXV (mimicry), XXVI (sense of god and religion), and XXVII (perseverance). We acknowledge that some of the associations may be less obvious. For example, the link between Faculty XII (sense of locality) and the lifestyle measure “Time spent doing light physical activity” is an assumption that physically active people are more likely to get out of the house. Although keeping in mind that the lifestyle measures have important clinical and scientific uses, all associations here were made in a spirit of *mirth.*

**Table 1.** Faculties and associated Biobank lifestyle measures.

| # | Faculty | Biobank lifestyle measure |
| --- | --- | --- |
| I | Impulse to propagation<br>(*Amativeness*) | Lifetime number of sexual partners |
| II | Tenderness for the offspring or parental love<br>(*Philoprogenitiveness*) | People in the house related to participant<br>(son/daughter/mother/father) |
| III | Friendly attachment or fidelity<br>(*Adhesiveness*) | People in the house not related to participant<br>(husband/wife/partner/other) |
| IV | Valour, self-defence<br>(*Combativeness*) | Solicitor, lawyer, barrister, judge (job) |
| V | Murder, carnivorousness<br>(*Destructiveness*) | Beef intake |
| VI | Sense of cunning<br>(*Cunning*) | Scientist (job) |
| VII | Larceny, sense of property<br>(*Acquisitiveness*) | Number of vehicles in household |
| VIII | Pride, arrogance, love of authority<br>(*Self-Esteem*) | Banker (job) |
| IX | Ambition and vanity<br>(*Love of Approbation*) | Financial situation satisfaction |
| X | Circumspection<br>(*Cautiousness*) | Alcohol intake frequency |
| XI | Aptness to receive an education or the memoria realis<br>(*Eventuality and Individuality*) | Age completed full time education |
| XII | Sense of locality<br>(*Locality*) | Time spent doing light physical activity |
| XIV | Words, verbal memory<br>(*Words*) | Letter fluency |
| XV | Faculty of language<br>(*Language*) | Authors, writers (job) |
| XVI | Disposition for colouring, delighting in colours<br>(*Colouring*) | Photographers, painter (job) |
| XVII | Sense for sounds, musical talent<br>(*Tune*) | Music profession (job) |
| XVIII | Arithmetic, counting, time<br>(*Number*) | Mathematician (job) |
| XIX | Mechanical skill<br>(*Constructiveness*) | Hand grip strength (right) |
| XX | Comparative perspicuity, sagacity<br>(*Comparison*) | Concept interpolation |
| XXI | Metaphysical perspicuity<br>(*Causality*) | Clergy (job) |
| XXII | Causality, sense of inference<br>(*Mirthfulness*) | Writer, actor, comedian (job) |
| XXIII | Poetic talent<br>(*Ideality*) | Poet (job) |
| XXIV | Good nature, compassion, moral sense<br>(*Benevolence*) | Charity (job) |

Figure 3 summarises the distribution of responses obtained for the lifestyle measures for the available subjects. The numbers of subjects sampled for each category were rather large, except for the “job”-based lifestyle measures (see Table 1). We ultimately left these faculties out of the final phrenological analysis, in which scalp morphology was correlated against personal measures.

**Figure 3:**
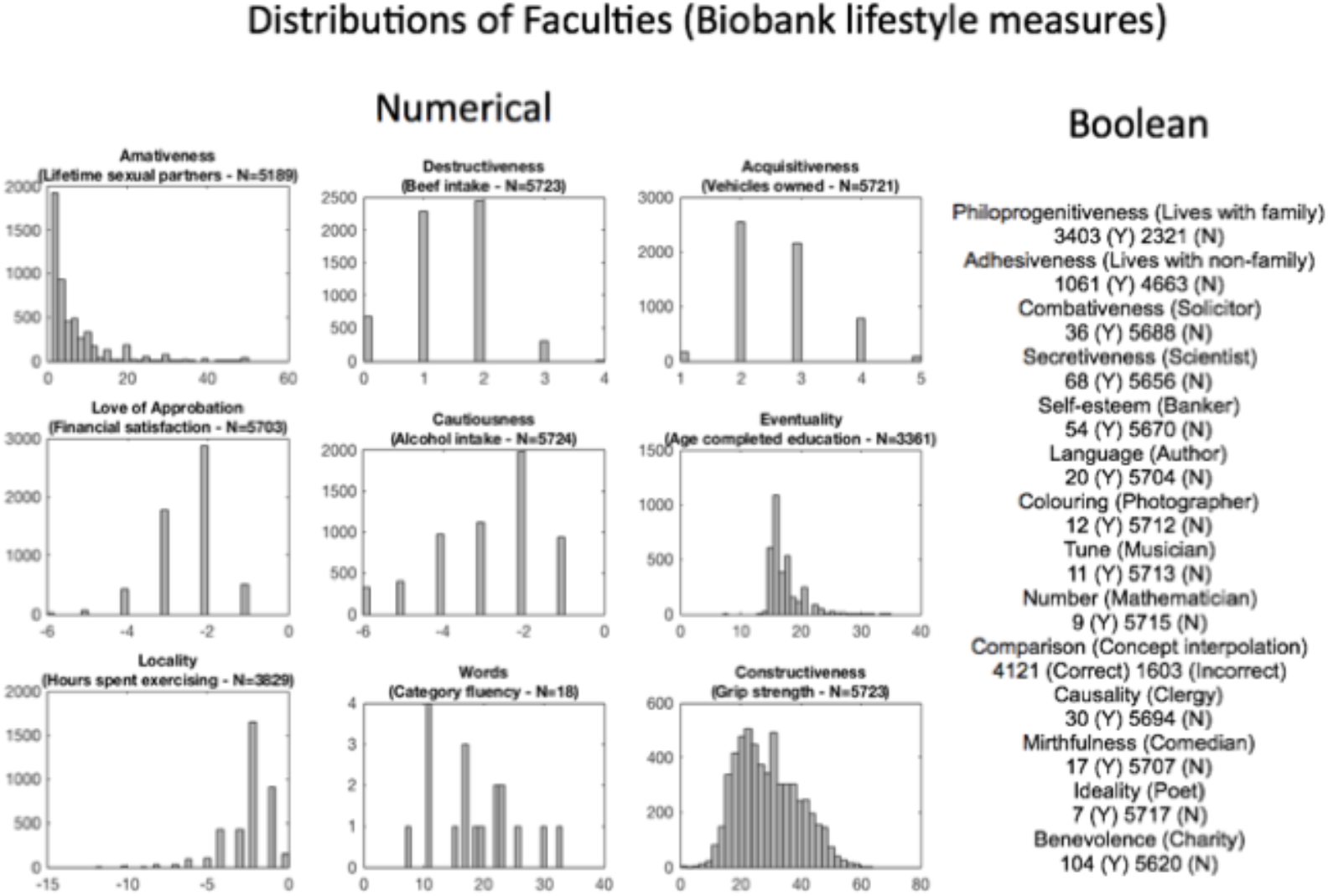
Distributions of faculties (Biobank measures). We matched Gall’s faculties against a set of personal measures that were acquired by UK Biobank. Three subject measures (financial satisfaction, alcohol intake, and time spent exercising) were multiplied by −1 to correlate positively with the corresponding faculties. Amativeness had a long tail (values going up to 1,000); although these were cut out in the figure, no values were excluded from the GLM analysis. For the numerical faculties, N refers to the total number of subjects; for the Boolean faculties, N refers to the number of subjects who answered “no”.

### Relating local scalp morphology to personal measures

Starting with the main claim of phrenology, that bumps on the head relate to individual traits, we used multivariate regression to search for associations between local scalp curvature and Biobank lifestyle measures. More concretely, we modelled vertex-wise scalp curvature against lifestyle measures including gender and age as nuisance regressors. For the binary measures, such as whether or not someone was a banker, the regression model could be set up as an unpaired t-test. For illustration purposes, all resulting t-statistics were converted to z-statistics (see Discussion). In total, we note that there were 14 binary lifestyle measures (Faculties II, III, IV, VI, VIII, XV, XVI, XVII, XVIII, XX, XXI, XXII, XXIII and XXIV) and nine non-binary measures (Faculties I, V, VII, IX, X, XI, XII, XIV, and XIX) all shown in Figure 3. However, because there were low numbers in some of the binary measures (Faculties IV, VI, VIII, XV, XVI, XVII, XVIII, XIX, XXI, XXII, XXIII, XXIV), these were omitted from the results.

We calculated vertex-wise p-values for the null hypothesis (that there should be no association between scalp curvature and lifestyle measures). In order to control for multiple comparisons across the scalp, which is something that phrenologists failed to the best of our knowledge to report, we used resel-based correction and Random Field Theory^26-28^ given a significance threshold of 0.05 (Bonferroni-corrected across the faculties tested).

### Relating local scalp morphology to local brain morphology

In order to test the second claim of phrenology, that bumps on the head should reflect the underlying shape of the cerebral cortex, we correlated each subject's local scalp curvature (described above) with a local index of brain gyrification (projected onto the scalp). This gyrification index was quantified using a surface ratio, corresponding to the amount of cortical surface packed within a limited spherical volume at every point on the cortex^29^. For data we extracted the cortical (pial) surface from each subject's T1-weighted scan using FreeSurfer^30^. In order to summarise the surface ratio of the cortex underlying each scalp vertex, we used the average surface ratio within a 20mm sphere centred around the nearest cortical vertex. Once both measures (scalp curvature and cortical convolution) were mapped onto the scalp surface, we could correlate the two measures and answer the question of whether scalp morphology may be considered a proxy for underlying brain morphology.

## Results

The phrenological analyses produced no statistically significant or meaningful effects.

## Discussion

The present study sought to test in the most exhaustive way currently possible the fundamental claim of phrenology: that measuring the contour of the head provides a reliable method for inferring mental capacities. We found no evidence for this claim. First, we explored the effect on local scalp curvature of underlying brain gyrification, given that phrenology assumes a relationship between head and brain morphology. We found that brain gyrification explains very little of the variance in local scalp curvature (Figure 4). Second, we correlated local scalp curvature with a set of lifestyle measures, interpreted as Victorian “faculties” (e.g. “lifetime number of sexual partners” was used as a proxy for the faculty of “Amativeness”, or the “impulse to propagation”). Despite the size of our sample and automation of our methods, we found no evidence to support phrenology's fundamental claim. The regions depicted on phrenological busts (Figure 1) therefore should not be trusted. According to our results, a more accurate phrenological bust should be left *blank* since no regions on the head correlate with any of the faculties that we tested. But *even below the level of statistical significance*, we found historic phrenological predictions to be uninsightful; for example, Figure 5 shows the unthresholded z-statistic map for correlations between local head curvature and lifetime number of sexual partners (“Amativeness”). Unsurprisingly, the “frontal horn” area that we point out does not correspond to ROIs proposed by phrenologists, which included areas at the *back* of the skull^25^. For the reckless, zealous or simply curious reader, we include the remaining unthresholded z-statistic maps (none statistically significant) in the Supplementary Materials (Supplementary Figure 2 to Supplementary Figure 23). We did not analyse the relationship between lifestyle measures and brain morphology, since many such relationships are known and uncontroversial within 21^st^ century neuroscience^31-33^. What is peculiar about phrenology is its emphasis on the outer head (i.e. skull and scalp) as an indirect measure of the brain, and thus of personality and behaviour.

**Figure 4:**
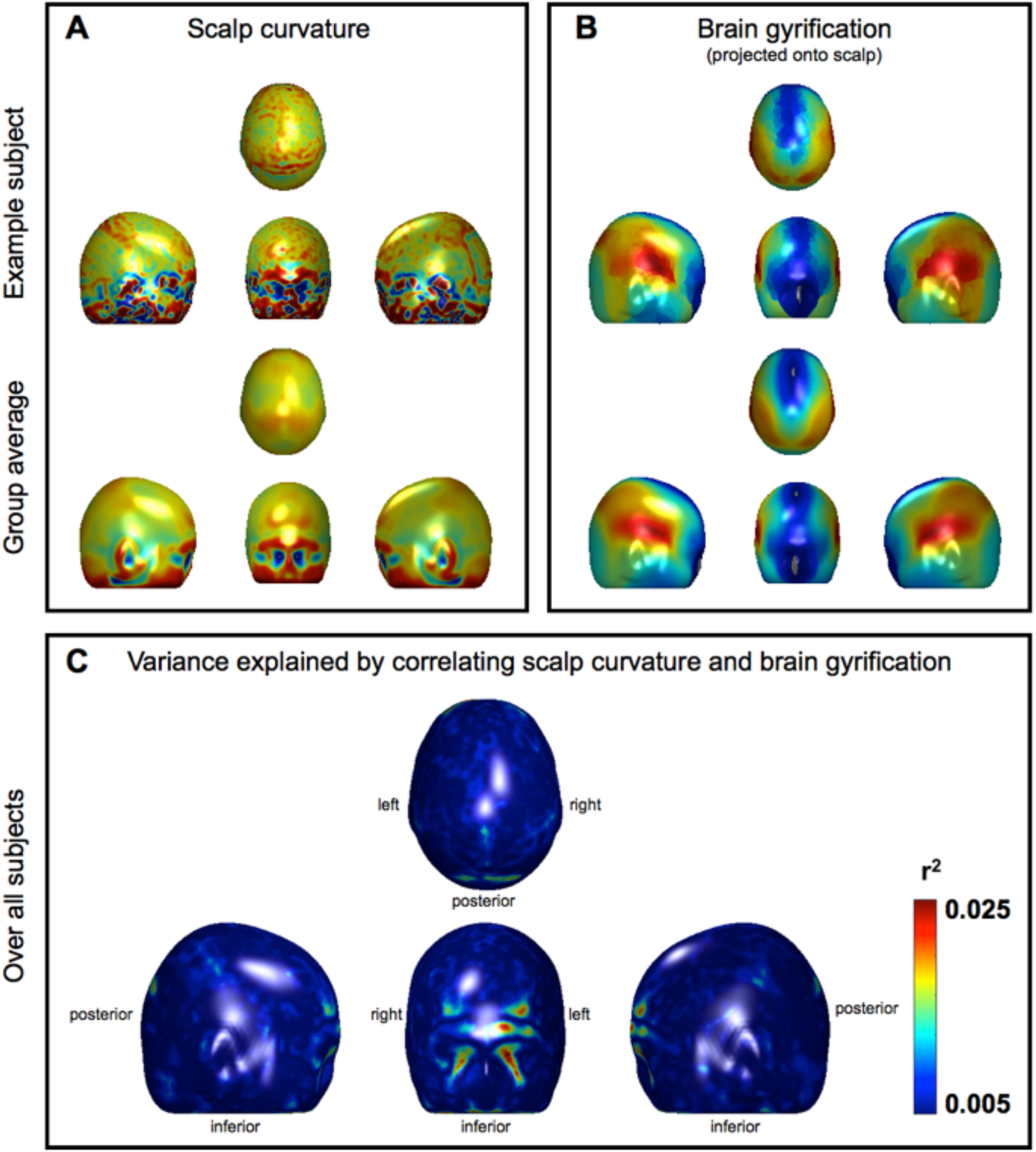
Scalp curvature, brain gyrification, and the variance explained by correlating the two. Panel A: Example scalp curvature data from a single subject (upper panel) and averaged over the entire cohort (lower panel). Red/Blue represents positive/negative (i.e. convex/concave) curvature values. Panel B: Example brain gyrification projected onto the scalp for a single subject (upper panel) and averaged over the entire cohort (lower panel). Red/Blue represents degree of gyrification (note large index values, in red, laterally over the Sylvian fissures). Panel C: Variance explained by correlating scalp curvature and brain gyrification. Note that the r2 values are very small; the “strongest” effects only explain about 0.025% of the variance (leaving 97.5% unexplained). The largest “effects” are also marginalised to the facial region, which is irrelevant to a great number of phrenological accounts and probably an artefact. All data have been projected onto the mean head surface.

**Figure 5:**
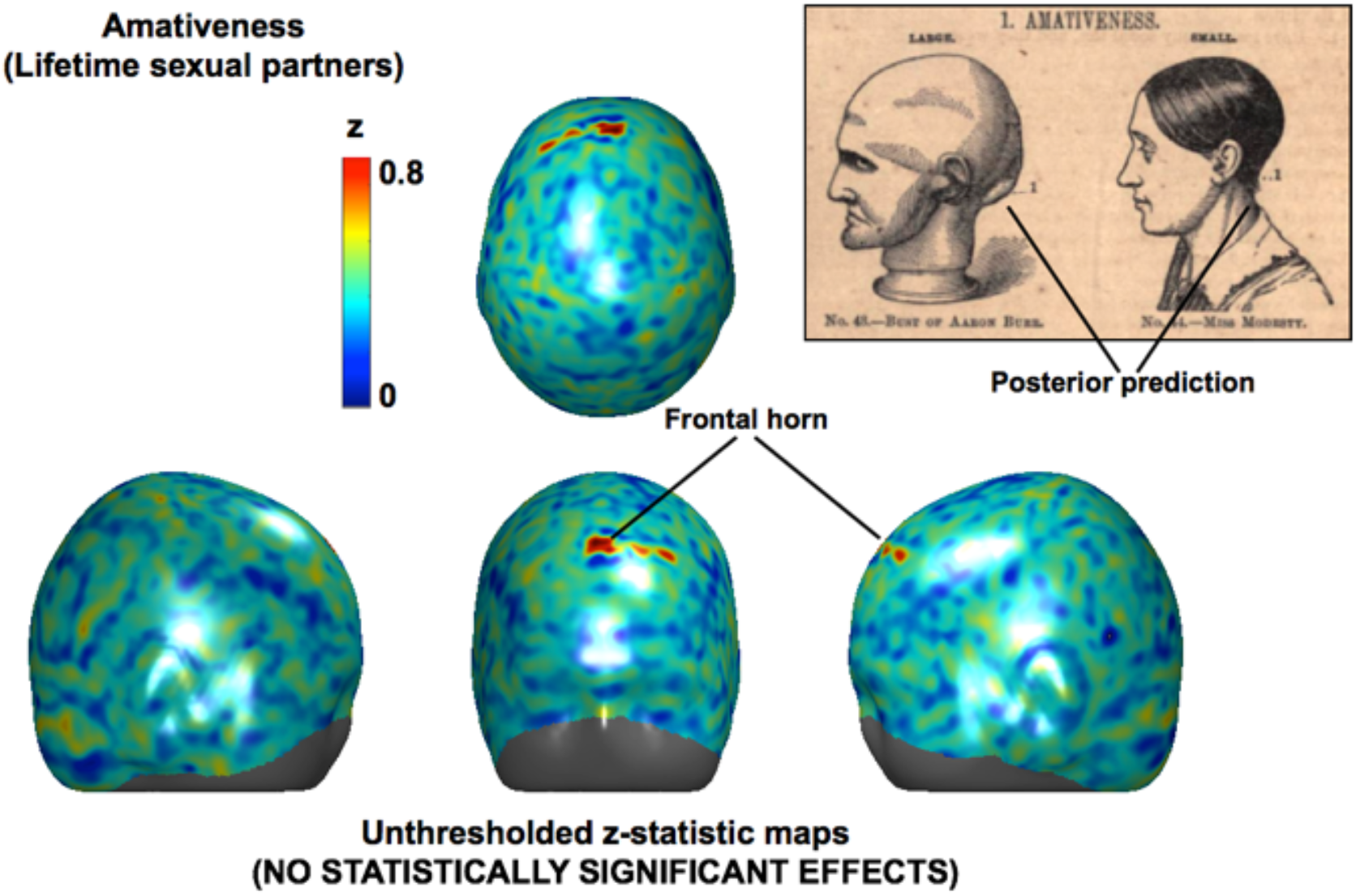
Illustration of over-interpreting null results. The scalp projections show an unthresholded z-statistic map of correlations between local scalp curvature and lifetime number of sexual partners, which has been overenthusiastically annotated with interpreted effects (i.e. the “frontal horn”). The results might be compared with those of the infamous “dead salmon” study, which highlighted the importance of correcting for multiple comparisons^34^. Please note that when thresholds for multiple comparisons were applied, none of the z-scores in this figure reached statistical significance. Also damning is the fact that the “frontal horn” area does not correspond to regions of interest predicted by 19^th^ century phrenologists. The upper-right panel depicts a prediction for “Amativeness” on the *opposite* side of the skull^25^.

The strengths of our approach are the automation of head measurements from MRI data and number of subjects studied. Because the analysis methods were automated, the number of subjects studied could easily number in the thousands. By contrast, although phrenologists had access to quantitative tools like the measuring tape and caliper, and some attempt was even made to automate the measuring procedure as evidenced by the psychograph (Figure 1), phrenology typically relied on “palpation” (the manual examination of subjects’ heads, which counted in the 19^th^ century as digital technology). Reference materials including phrenology charts and Fowler heads (Figure 1) were the results of underpowered studies, including perhaps only an anecdotal handful subjects^24^.

Set against the strengths of our study, an apparent weakness is our use of 19^th^ century “faculty psychology” with its description of human nature in idiosyncratic terms like “Amativeness” and “Philoprogenitiveness”^22^, and grouping together of attributes like “eats meat” and “likes to kill”, which may strike us today as odd^24^. In other words, it might be objected that we should have used a more recent *ontology.* However, phrenology's “faculty psychology” is not as different from current ontologies as it first seems. One may readily find examples of the same faculties in the neuroimaging literature, albeit under different names (Table 2). We were also interested in grounding our study in Victorian concepts, despite an emphasis on 21^st^ century methods. The lifestyle features that we selected also ranged over a wide number of behavioural and cognitive domains (e.g. motor skills, language, spatial awareness, decision making, etc.), so regardless of ontology we hope to have covered many topics of interest.

**Table 2.** examples of nineteenth-century phrenology faculties in modern neuroimaging studies (of the brain). Adapted from Poldrack^22^.

| **Phrenological faculty** | **Modern neuroimaging equivalent** | **Associated regions** | **References** |
| --- | --- | --- | --- |
| Impulse to propagation<br>(*Amativeness*) | Viewing of romantic lover vs. other individuals | Basal ganglia | Aron et al.<sup>35</sup> |
| Ambition and vanity<br>(*Love of Approbation*) | Activation for judgement about self vs. others | Medial prefrontal cortex | Ochsner et al.<sup>36</sup> |
| Circumspection<br>(*Cautiousness*) | Activation correlated with harm avoidance | Nucleus accumbens | Matthews et al.<sup>37</sup> |
| Arithmetic, counting, time<br>(*Number*) | Activity correlated with arithmetic skill | Angular gyrus | Menon et al.<sup>38</sup> |

As to the objection that phrenology was already a known dead-end scientifically, and that its claims did not need to be tested rigorously, it is indeed hard to find a time in history when phrenology was not seriously criticised. Even in 1815, the year that Spurzheim published his influential book on Gall's method^1^, phrenology was dismissed by one reviewer as “a piece of thorough quackery from beginning to end”^39^. Not only did the reviewer take issue with the use of palpation as an indirect method for measuring the brain and its mental faculties, but he also objected to the idea the brain might be composed of multiple specialised components, writing^39^ (p. 243):

> “The cases in which portions of various sizes have been removed from almost all regions of this organ [the brain], without the slightest affection either of Intellect or Inclination, are numerous and most unequivocal.”

Despite the reviewer's objection, this second idea, known today as “functional specialisation” or “segregation”^40^, has proven of central practicality to our understanding of the brain since the first of Broca's famous case studies^4^. This highlights the importance of empiricism and of testing improbable sounding theories. We would argue that phrenology's first idea, that the shape of the head might reflect brain function, is not *a priori* incoherent. It is certainly true that the shape of the head reflects mental capacities in extreme pathological cases^41^, such as hydrocephalus, where increasing head size could reflect progressive ventriculomegaly. Even in the healthy population, adequate childhood nutrition might result both in increased intelligence scores and in parallel skull growth, such that one might detect a correlation between intelligence tests and local scalp curvature. The possibility of this outcome shows that the scalp-curvature hypothesis could not be refuted by armchair methods alone, but required empirical testing. We of course acknowledge that science cannot test all hypotheses, but rather that, because of limited resources, scientists much choose between experiments^42^. Therefore it would not have been realistic, or perhaps even ethical, to acquire MRI for thousands of subjects with the purpose of testing a long-abandoned theory. However, one of many benefits that big data projects like the UK Biobank confer is that they provide resources for answering questions that might otherwise have remained untested, or even untestable.

In closing, we hope to have argued convincingly against the idea that local scalp curvature can be used to infer brain function in the healthy population. Given the thoroughness of our tests, it is unlikely that more scalp data would yield significant effects. It is true that further work might focus on the *inner* (rather than outer) curvature of the skull, perhaps formalising a virtual method for creating endocasts^43,44^. In any case, we would advocate that future studies focus on the *brain.* Although written in a light-hearted spirit, our study demonstrates the feasibility of applying to cranial data the standard methods of neuroimaging (like registration, normalisation, random field theory, and mass univariate analysis). One potential application of these methods would be the clinical treatment of craniosynostosis. In extreme cases of craniosynostosis, paediatric surgeons will separate the fused bones in a baby's head to increase the size of the cranial vault, thereby creating space for brain growth. However, because of the inherent risks of surgery, there are many “border” cases that are not operated on, where the use of neuroimaging methods on the skull could be used to track (or maybe retrospectively evaluate) correlations between local head shape and cognitive development. This would be useful for evaluating whether developmental impairments should motivate that similar “border” cases be operated on in future, and, incidentally, is closer to the original serious scientific and clinical motivation behind acquiring the UK Biobank.

## Acknowledgements

This research has been conducted using the UK Biobank Resource under Application Number 8107.

## Contributors

SJ and OPJ conceived of the study over pints at our local pub, the White Hart; all authors contributed to data analysis and writing.

## Ethical approval

Not required.

## Transparency statement

The lead author (OPJ) affirms that the manuscript is an honest, accurate, and transparent account of the study being reported; that no important aspects of the study have been omitted; and that any discrepancies from the study as planned have been explained.

## Data sharing

The data are available online (http://www.ukbiobank.ac.uk/).

